# *In Vivo* Delivery of Therapeutic Molecules by Transplantation of Genome-Edited Induced Pluripotent Stem Cells

**DOI:** 10.1101/2023.01.03.522057

**Authors:** Ittetsu Nakajima, Takahiro Tsukimura, Terumi Ono, Tomoko Shiga, Hiroshi Shitara, Tadayasu Togawa, Hitoshi Sakuraba, Yuichiro Miyaoka

## Abstract

Human induced pluripotent stem cells (iPSCs) have already been used in transplantation therapies. Currently, cells from healthy people are transplanted into patients with diseases. With the rapid evolution of genome editing technology, genetic modification could be applied to enhance the therapeutic effects of iPSCs, such as the introduction of secreted molecules to make the cells a drug delivery system. Here, we addressed this possibility by utilizing a Fabry disease mouse model, as a proof of concept. Fabry disease is caused by the lack of α-Galactosidase A (GLA). We previously developed an immunotolerant therapeutic molecule, modified α-N-acetylgalactosaminidase (mNAGA). We confirmed that secreted mNAGA from genome-edited iPSCs compensated for the GLA activity in GLA-deficient cells using an *in vitro* co-culture system. Moreover, iPSCs transplanted into Fabry model mice secreted mNAGA and supplied GLA activity to the liver. This study demonstrates the great potential of genome-edited iPSCs secreting therapeutic molecules.

## Introduction

Induced pluripotent stem cells (iPSCs) have great potential as resources for cell therapy (Takahashi et al., 2007). Since the first clinical trial of iPSC-based transplantation into an age-related macular degeneration patient in 2014 (Mandai et al., 2017), several transplantation therapies using iPSC-derived platelets (jRCTa050190117), corneal epithelial cell sheets (jRCTa050190084), cardiomyocyte sheets (jRCT2053190081), and dopaminergic progenitors (UMIN000033564) have proceeded to clinical trials.

So far, these transplantations have utilized unmodified iPSCs from healthy patients (Xu et al., 2019). However, we can potentially improve the therapeutic effects of iPSCs by genome editing as long as the safety is guaranteed (Urnov et al., 2010) (Joung and Sander, 2013) (Jinek et al., 2012). For example, disruption of some of the HLA genes can expand the range of people who can receive the cells without immune responses (Xu et al., 2019). Genomic insertion of chimeric antigen receptor (CAR) has been utilized for T-cell immunotherapy (Iriguchi et al., 2021). Another prospect of genetic modification is to make iPSCs that secrete therapeutic molecules. Recently, the implantation of genome-engineered iPSCs that secrete interleukin-1 (IL-1) in rheumatoid arthritis mice demonstrated the potential of this strategy (Choi et al., 2021). However, the transplantation of iPSCs to deliver therapeutic molecules *in vivo* has not been established as a therapeutic option. Therefore, we decided to test whether iPSCs with genetic modifications can deliver therapeutic molecules *in vivo* in mice by focusing on Fabry disease as a proof of concept.

Fabry disease (OMIM 301500) is an X-linked genetic disorder (Fabry, 1898) that is caused by a deficiency of α-Galactosidase A (GLA, EC 3. 2. 1. 22). GLA is a lysosome enzyme that is partially secreted, and incorporated by other cells. Therefore, the loss of GLA results in accumulation of its substrates such as globotriaosylceramide (Gb3) and globotriaosylsphingosine (Lyso-Gb3) (Sweeley and Klionsky, 1963). The accumulation leads to various symptoms, including cardiomyopathy, renal failure, and stroke (Germain, 2010).

Currently, there are two major treatments for Fabry disease: enzyme replacement therapy (ERT) (Eng et al., 2001) (Schiffmann et al., 2001) and pharmacological chaperone therapy (PCT) (Germain et al., 2016). In ERT, the recombinant human GLA is intravenously infused every two weeks. However, this infusion can lead to the development of antibodies against GLA and hampers the therapeutic effect in some patients (Concolino et al., 2018). In PCT, small molecules that stabilize GLA are orally administered. However, PCT has no effect in patients with the complete loss of GLA (Benjamin et al., 2017) (Kobayashi et al., 2019).

To solve these problems, we aimed to establish a new iPSC transplantation therapy for Fabry disease using a modified enzyme that we previously developed, modified α-N-acetylgalactosaminidase (mNAGA). mNAGA was created by altering the substrate specificity of NAGA, which is a paralog of GLA, into that of GLA (Tajima et al., 2009). Because mNAGA maintains the original antigenicity, this modified enzyme has no immunological cross-reactivity with GLA, while having the GLA enzymatic activity. In this study, we tested if transplantation of iPSCs secreting mNAGA by genome editing could supply the GLA activity *in vivo*. Our strategy of cell therapy using genome-edited iPSCs can be applied to various diseases caused by loss of functional molecules.

## Results

### Generation of human iPSCs secreting mNAGA by genome editing

First, we established a human iPSC line that secretes mNAGA to test its therapeutic effects in Fabry model mice. To exclude the possible immunogenic reactions caused by the endogenous GLA of iPSCs in patients, we knocked-down *GLA* by introducing a 6-bp deletion including the start codon by CRISPR-Cas9 (Figure 1a) (hereafter called GLA-KO iPSCs). We confirmed the loss of GLA activity in GLA-KO iPSCs (Figure 1b).

**Figure 1.**
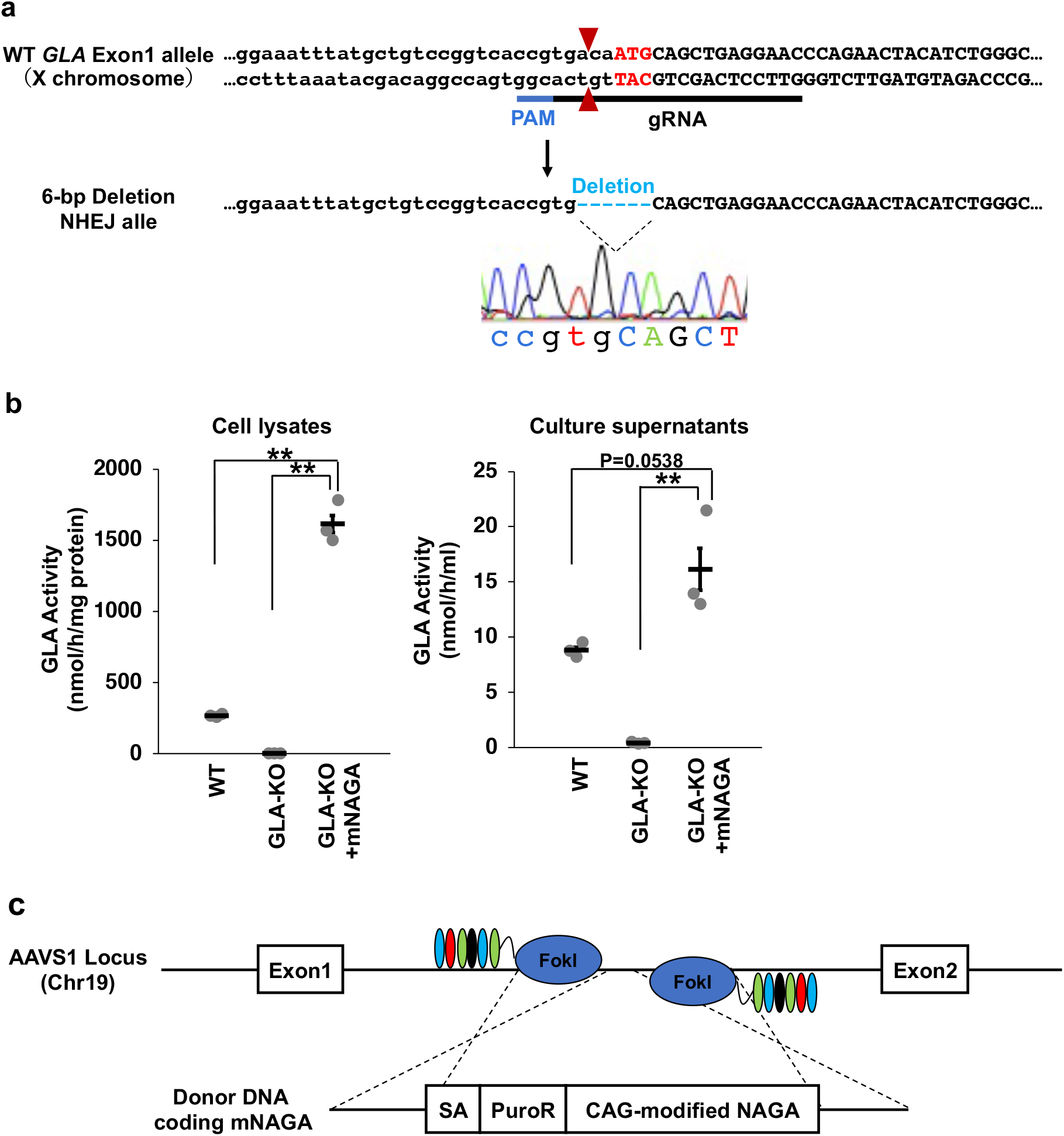
Establishment of mNAGA-secreting iPSCs. (a) Design of the disruption of the GLA gene in human iPSCs by CRISPR-Cas9. We introduced a 6-bp deletion containing the start codon. The cut site and start codon are shown by red triangles and red letters, respectively. Sangar sequencing data around the deletion are shown. (b) GLA activity in the lysates or culture supernatants of wild-type, GLA-KO, and GLA-KO+mNAGA iPSCs (n=3). *P<0.05 and **P<0.01. (c) Design of knock-in of mNAGA cDNA driven by the CAG promoter into AAVS1 using TALENs.

Next, to establish an iPSC line stably expressing mNAGA, we heterozygously knocked-in mNAGA cDNA driven by the CAG promoter into adeno-associated virus integration site 1 (AAVS1), a safe harbor locus, by TALENs (Figure 1c) (hereafter called GLA-KO+mNAGA iPSCs). This cell line exhibited higher GLA activity than wildtype (WT) and GLA-KO iPSCs both in cell lysate and in culture supernatant (Figure 1b). Thus, we successfully generated iPSCs for a potential cell therapy for Fabry disease by genome editing.

### mNAGA secreted from iPSCs was taken up by GLA-deficient cells *in vitro*

For cell therapy, mNAGA secreted from transplanted iPSCs must be taken up by patients’ cells. Therefore, we investigated whether GLA-deficient cells could pick up mNAGA secreted from iPSCs by co-culture using transwells (Figure 2a).

**Figure 2.**
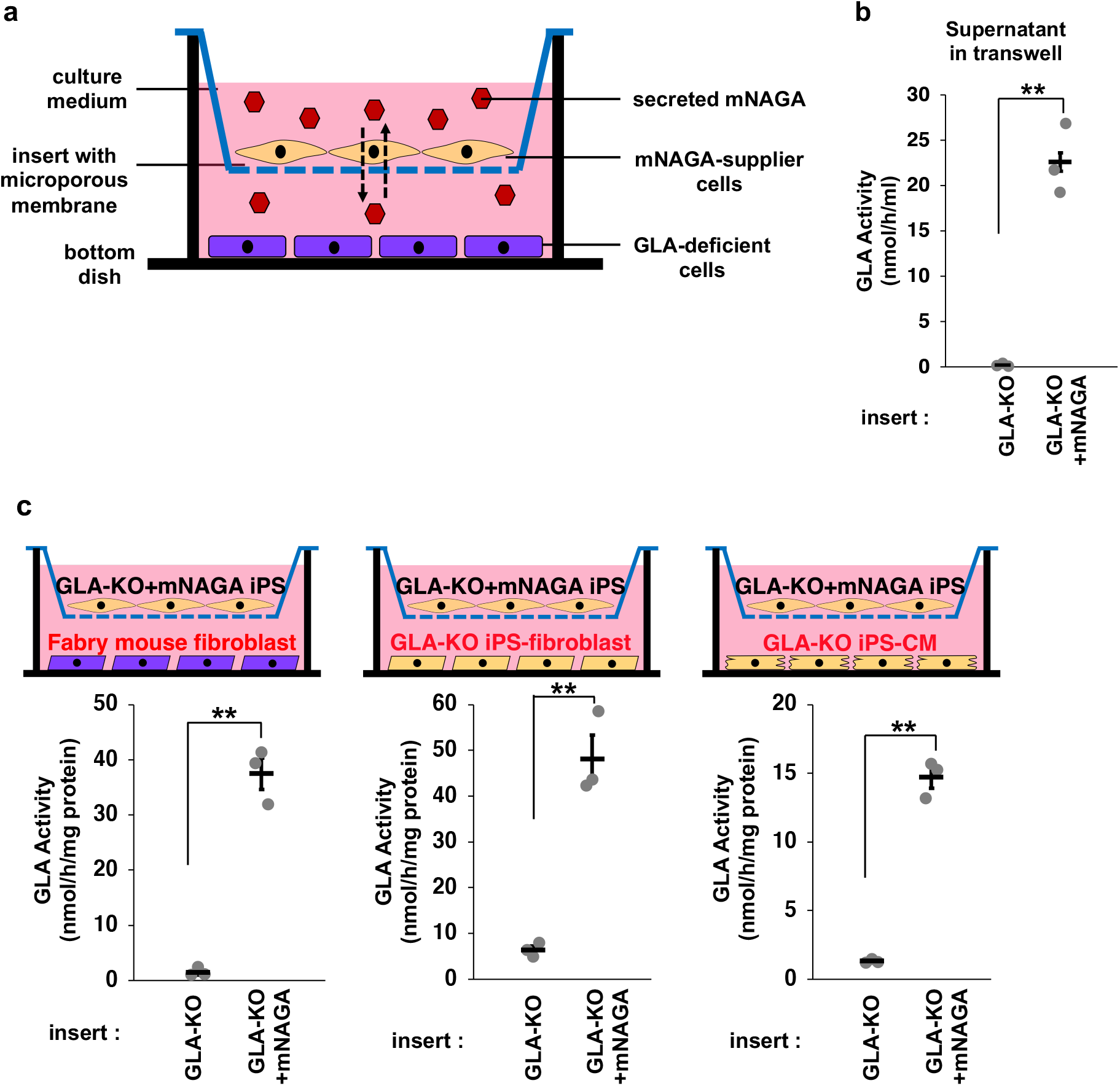
*In vitro* delivery of mNAGA secreted from iPSCs to GLA-deficient cells. (a) Schematic illustration of *in vitro* delivery of mNAGA using transwells. (b) GLA activity of the culture supernatants of GLA-KO or GLA-KO+mNAGA iPSCs (n=3). *P<0.05 and **P<0.01. (c) GLA activity of mouse fibroblast lysates co-cultured with GLA-KO or GLA-KO+mNAGA iPSCs (n=3). *P<0.05 and **P<0.01. (d, e) GLA activity of iPSC-derived fibroblasts (d) and iPSC-derived cardiomyocytes (e) co-cultured with GLA-KO or GLA-KO+mNAGA iPSCs (n=3). *P<0.05 and **P<0.01.

First, we investigated whether mNAGA secreted from iPSCs can be incorporated into fibroblasts derived from Fabry mice without functional GLA, as described previously (Ohshima et al., 1997). We co-cultured Fabry mouse fibroblasts on the bottom of a transwell and undifferentiated GLA-KO or GLA-KO+mNAGA iPSCs on the insert for two days. We confirmed the secretion of mNAGA from the iPSCs by measuring the GLA activity in the culture medium (Figure 2b). As expected, the GLA activity in the co-cultured Fabry mouse fibroblasts was higher than in the control. These findings indicated that secreted mNAGA restored the GLA activity of Fabry mouse fibroblasts (Figure 2c).

Next, to simulate situations more relevant to actual cell therapy, we co-cultured GLA-KO iPSC-derived fibroblasts or cardiomyocytes with GLA-KO+mNAGA iPSCs, and observed the restoration of the GLA activity in these cells (Figure 2d, e). These results demonstrated that mNAGA secreted from GLA-KO+mNAGA iPSCs can be incorporated into mouse and human GLA-deficient cells and complement the lost GLA activity.

### Generation of an immunodeficient Fabry mouse model

Next, we generated an immunodeficient Fabry mouse model in which human cells can be transplanted and engrafted. We disrupted the *Gla* gene in NOD.CB17-*Prkdc*^*scid*^/J (NOD SCID) mice, a mouse strain without functional T and B cells (Shultz et al., 2007) by injecting Cas9 mRNA and sgRNA targeting exon1 of mouse *Gla* (Figure 3a). We obtained 25 founder mice with various insertions and/or deletions out of 100 injected zygotes (Table S1). Among them, we established one mouse line with a 28-bp deletion (*Gla*^Δ28^) spanning the start codon of *Gla* (Figure 3a). We confirmed that the GLA activity was lost in the liver, heart, and kidney of this *Gla*^Δ28^ mouse strain (Figure 3b).

**Figure 3.**
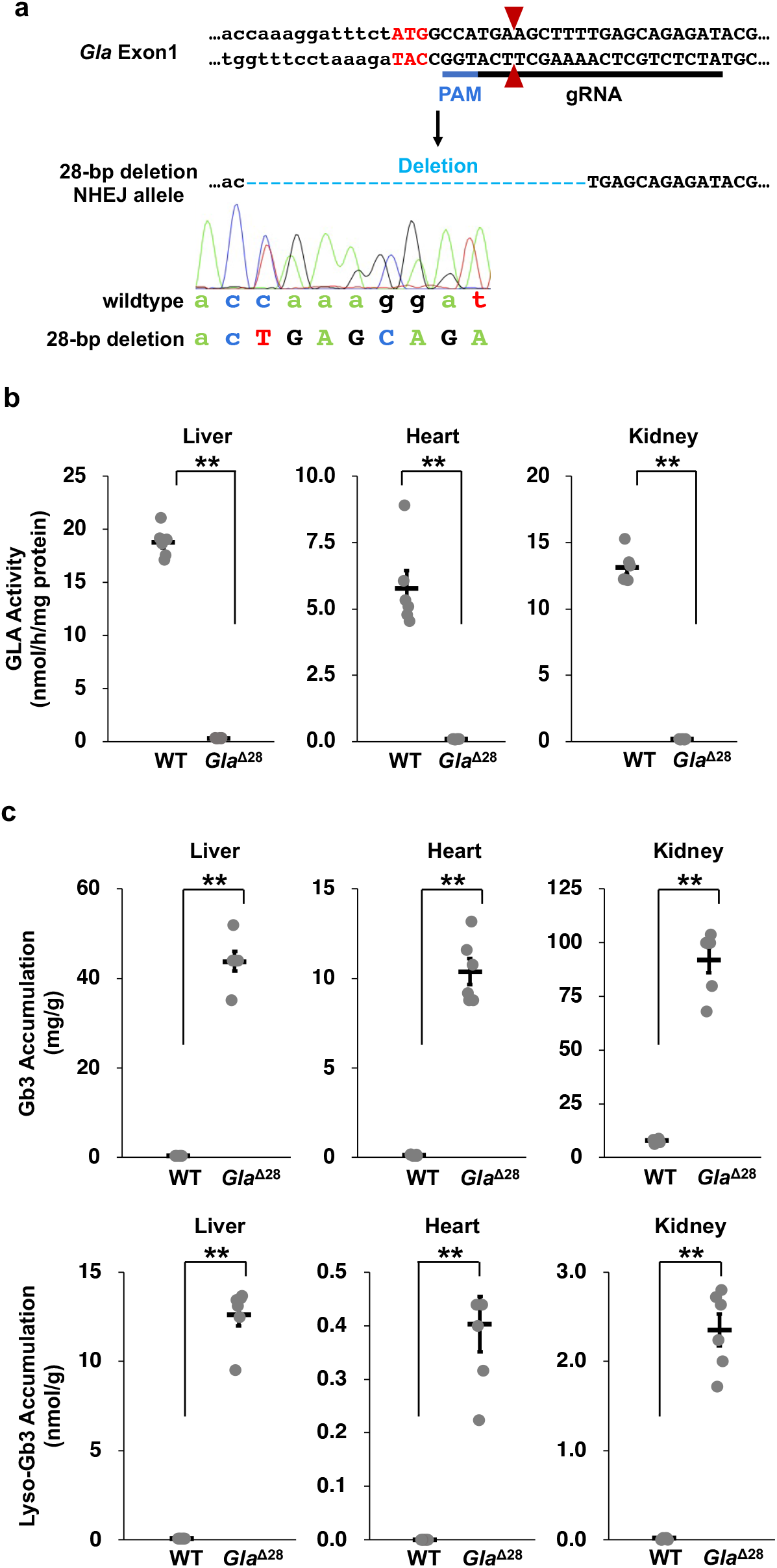
Generation of the immunodeficient Fabry mouse model. (a) Design of the disruption of *Gla* in NOD SCID mouse zygotes. The cut site and the start codon are shown by red triangles and red letters, respectively. The allele with a 28-bp deletion containing the start codon identified as a null allele, *Gla*^Δ28^, is shown. The Sanger sequencing data show the wild-type allele and the 28-bp deletion allele in a heterozygous mouse. (b) The GLA activity in the liver, heart, and kidneys of male wild-type and *Gla*^Δ28^ mice (n=6). *P<0.05 and **P<0.01. (c) Accumulation of Gb3 and Lyso-Gb3, substrates of GLA, in the liver, heart, and kidneys of male wild-type and *Gla*^Δ28^ mice (n=6). *P<0.05 and **P<0.01.

Next, we measured the amounts of Gb3 and Lyso-Gb3, substrates of GLA, in the *Gla*^Δ28^ mouse strain by tandem mass spectrometry. As we expected, these substrates were accumulated in all analyzed tissues (Figure 3c). These results confirmed that the NOD SCID *Gla*^Δ28^ mouse strain we established could serve as an immunodeficient Fabry mouse model.

### Transplantation of GLA-KO+mNAGA iPSCs to NOD SCID Fabry mice

Finally, we transplanted GLA-KO+mNAGA iPSCs into the NOD SCID Fabry mice. As the most efficient strategy to transplant the largest number of human cells, we formed teratomas in these mice.

We injected 1.0×10^6^ undifferentiated GLA-KO or GLA-KO+mNAGA iPSCs into the testes of 7 or 8-week-old male NOD SCID *Gla*^Δ28^ mice. After 7 or 8 weeks, we observed teratomas in all treated mice (Figure 4a). We confirmed that GLA-KO+mNAGA iPSC-derived teratoma maintained the GLA activity (Figure 4b).

**Figure 4.**
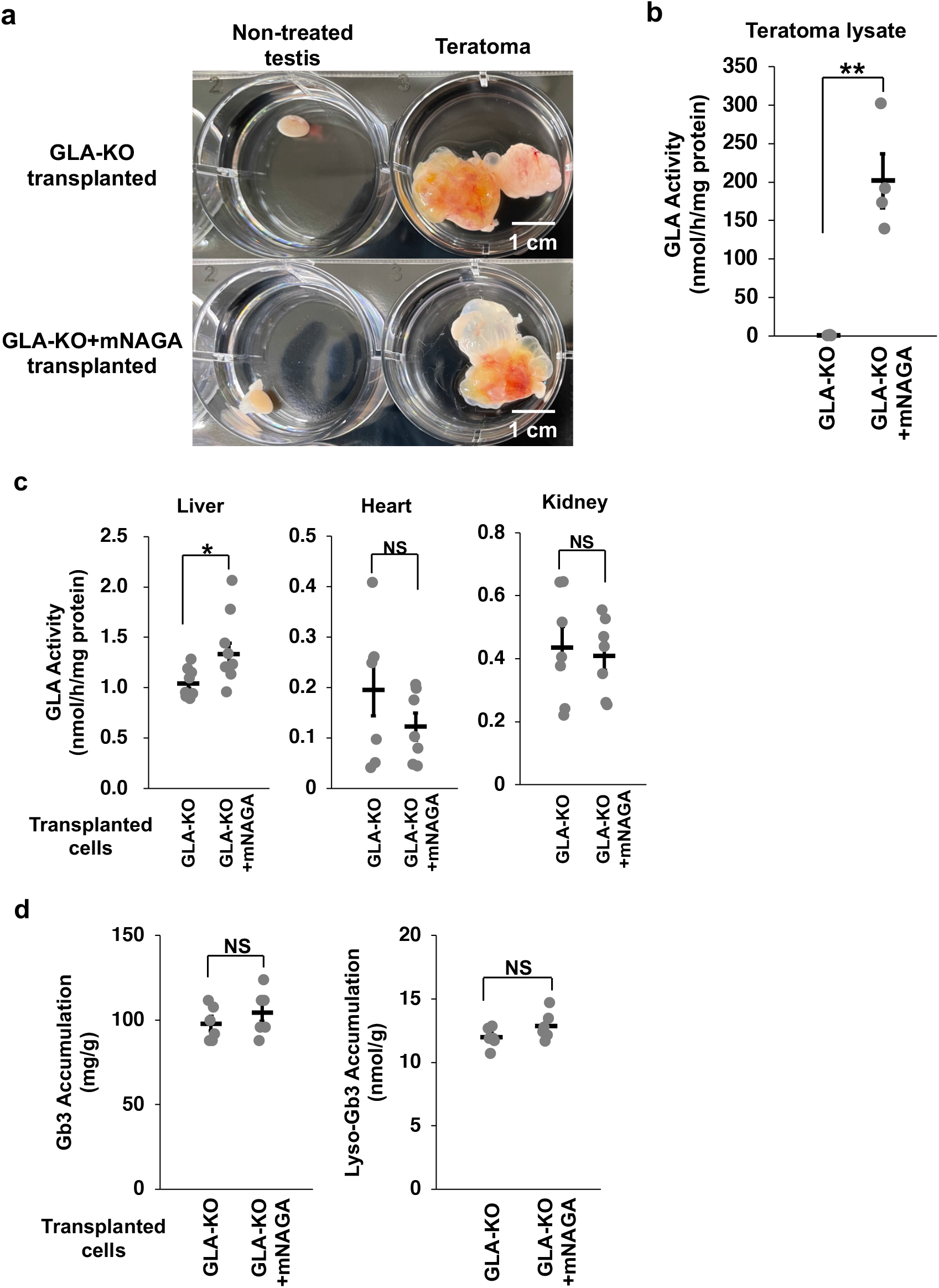
Transplantation of GLA-KO+mNAGA iPSCs to NOD SCID Fabry model mice. (a) Images of a non-treated testis and a testis with teratoma. Scale bars indicate 1 cm. (b) GLA activity in teratoma lysates derived from GLA-KO or GLA-KO+mNAGA iPSCs (n=4). *P<0.05 and **P<0.01. (c) GLA activity in the liver, heart, and kidneys of NOD SCID Fabry mice transplanted with GLA-KO or GLA-KO+mNAGA iPSCs (n=9 in only liver. n=6 in heart, and kidneys). *P<0.05, **P<0.01 and NS: not significantly different (P>0.1). (d) Amounts of Gb3 and Lyso-Gb3 in the liver of Fabry mice transplanted with GLA-KO or GLA-KO+mNAGA iPSCs (n=6). *P<0.05, **P<0.01 and NS: not significant (P>0.1).

To investigate whether transplanted GLA-KO+mNAGA iPSCs improved the GLA activity in these mice, we measured the GLA activity in the liver, heart, and kidneys. We observed no recovery of the GLA activity in the heart or kidneys. However, the GLA activity in the liver was significantly improved by the transplantation of GLA-KO+mNAGA iPSCs (P=0.0213) (Figure 4c). We also quantified the amounts of Gb3 and Lyso-Gb3 in the liver, but there was no detectable reduction of the substrates (Figure 4d). Taken together, these data indicate that although further optimization is required, we successfully delivered mNAGA from transplanted iPSCs to the liver *in vivo*.

## Discussion

We generated human iPSCs expressing and secreting mNAGA by genome editing. Furthermore, the GLA gene was disrupted to exclude the potential immunogenicity of the endogenous GLA in iPSCs. We think these genome editing strategies can be applied to other disorders associated with the loss of functional molecules.

We successfully demonstrated that mNAGA secreted from iPSCs could restore the GLA activity in GLA-deficient cells by co-culture experiments. We also successfully established an immunodeficient Fabry mouse model. Then, we transplanted iPSCs into these mice to form teratomas. Although teratoma formation is not applicable as a treatment option, it was the best experimental approach, because teratoma formation can engraft a large number of cells and the transplanted cells can be easily identified in the mouse body. One of the organs most affected by Fabry disease is the heart. Therefore, one promising therapeutic option is to transplant cardiomyocyte-sheets generated from iPSCs secreting mNAGA. GLA-KO+mNAGA iPSC-derived teratomas improved the GLA activity only in the liver of the treated Fabry model mice.

This result is consistent with a previous study reporting that the majority of recombinant GLA administered to mice is incorporated into the liver (Sakuraba et al., 2006). Despite the improved GLA activity, the liver levels of Gb3 and Lyso-Gb3 did not change. The liver GLA activity in mice treated with GLA-KO+mNAGA iPSCs was 1.33 nmol/h/mg protein, which was still much lower than in the WT mouse liver (18.80 nmol/h/mg protein). Therefore, we should enhance the production and secretion of mNAGA from iPSCs. The GLA activity in the GLA-KO+mNAGA iPSCs lysate was 6 times higher than that of WT. However, the difference in the culture medium was only 1.8 times. These results suggest that most mNAGA was retained in the lysosomes, and actively secreted (Medin et al., 1996). Several genes are involved in regulating the transportation of lysosomal enzymes (Yogalingam et al., 2008) (Medina et al., 2011) (van Meel et al., 2016). Thus, overexpression or disruption of these genes potentially enhances the secretion or uptake of mNAGA. Recently, it was reported that transplantation of engineered spheroids composed of mouse embryonic fibroblasts reduced the amount of Lyso-Gb3 in Fabry model mice (Kami et al., 2020). Therefore, the comparison of the two strategies would be beneficial.

In summary, this study demonstrated that genome edited iPSCs secreting therapeutic molecules could serve as not only a resource of cell transplantation but also a drug delivery system.

### Experimental procedures Genome editing of human iPSCs

For disruption of GLA, we used the P3 Primary Cell Kit and Nucleofector 4D device (Lonza). Two hundred thousand WTC11 iPSCs were transfected with 500 ng of pX330 with a gRNA targeting the first exon of the GLA gene (5’-GTTCCTCAGCTGCATTGTCA-3’) by the DS-138 program. Induced indels were detected by Droplet Digital PCR, and clones with indels were isolated by sib-selection (repeated limited dilutions), as described previously (Miyaoka et al., 2016).

For knock-in of mNAGA cDNA driven by the CAG promoter, we used the Human Stem Cell Nucleofector Kit-1 and Nucleofector 2b Device (Lonza). Two million iPSCs were transfected with 2 µg of each AAVS1-TALEN-L/R (Addgene #59025 and #59026) (González et al., 2014) and 5 µg of AAV-CAGGS-mNAGA donor plasmid. AAV-CAGGS-mNAGA was constructed by replacing the EGFP cDNA of AAV-CAGGS-EGFP (Addgene #22212) (Hockemeyer et al., 2009) with mNAGA cDNA. After transfection using Nucleofector 2b using the A-23 program, the treated cells were resuspended in mTeSR Plus with Y-27632 and plated into a Matrigel-coated 96-well plate. Two days after transfection, the medium was exchanged with mTeSR Plus with Y-27632 and 0.5 µg/ml Puromycin for selection of transfected cells. The cells were cloned by limiting dilution. We extracted genomic DNA from cloned cells and confirmed the knock-in by junction PCR and then Sanger sequencing.

### Measurement of the GLA activity using an artificial fluorogenic substrate

Hydrolysis of an artificial GLA substrate, 4-methylumbelliferyl-α-D-galactopyranoside (4MU-Gal), releases the fluorescent product, 4-MU (Mayes et al., 1981). Therefore, quantification of the amount of produced 4-MU can measure GLA activity. We mixed 40 µl of 4MU-Gal with 10 µl of tissue, culture supernatant, or cell lysate and incubated at 37°C for 30 min. The reaction was terminated by adding 950 µl of 0.2 M Glycine buffer pH10.7. We transferred 200 µl of the solution into a well of a 96 well black microplate (Greiner Bio-One). The fluorescence at 415-445 nm of 4-MU was measured by excitation at 365 nm using GloMax (Promega). For the GLA activity of tissue or cell lysates, the GLA activity per volume (nmol/h/ml) was converted to the activity per protein amount (nmol/h/mg) using the protein concentration measured by a Micro BCA assay.

### The mass spectrometric analysis of Gb3 and Lyso-Gb3

For measurement of Gb3 and Lys-Gb3 in organs and tissues of mice, a liquid chromatography (LC)-tandem mass spectrometry (MS/MS) analysis was performed according to a previous report (Tsukimura et al., 2021). Briefly, the organs and tissues were homogenized in MES buffer, and a 10 µl aliquot of the homogenate was mixed with a 70 µl aliquot of chloroform: methanol (1:2). Then a 10 µl aliquot of 5 µg/ml Gb3 (C17:0) (Matreya, LLC) and a 10 µl aliquot of 500 nmol/l stable isotope-labelled (one 13C and three deuteriums) Lyso-Gb3 (Nard Institute) were added as internal standards. The mixture was centrifuged and the supernatant was transferred to an LC vial. For LC, a Union UK-C8 column (20 × 3 mm ID., 3 µm; Imtakt Co.) was used, and the column oven was set at 30°C. Chromatographic separation was performed with a binary gradient consisting of a mobile phase of water containing 0.1% acetic acid and 2 mmol/l ammonium acetate, and methanol containing 0.1% acetic acid and 2 mmol/l ammonium acetate. The flow rate was 0.25 ml/min and the injection volume was 2 μl. Then, Gb3 isoforms and Lyso-Gb3 in the samples were detected by MS/MS using a LCMS-8040 triple quadrupole mass spectrometer (Shimadzu) equipped with an electrospray ionization interface in positive-ion mode. The multiple reaction monitoring (MRM) conditions were optimized with an automatic MRM optimization function. The calculations for measurement of Gb3 isoforms, and Lyso-Gb3 were performed using LabSolutions (Shimadzu), and the Gb3 contents of organs and tissues were calculated from the sums of those of the Gb3 isoforms.

### Co-culture of two types of cells using trasnswells

We used the bottom dish and the insert for GLA-deficient cells and mNAGA-supplier cells, respectively (Figure 2). These cells were plated in the bottom dish or the insert and separately cultured until the cells reached 100% confluence. Then, the insert was moved to the bottom dish for co-culture. For undifferentiated iPSCs, both the bottom dish and the insert were coated with Matrigel. We co-cultured for 3 days for mouse fibroblasts, and for 1 week for other cell types. We chose the medium for co-culture depending on the GLA-deficient cell types in the bottom dish.

### Transplantation of human iPSCs into mouse testes

iPSCs were detached using 500 µl of 0.5 mM EDTA (Nacalai Tesque) in PBS per well of a 6-well plate and resuspended in 1.5 ml of mTeSR Plus with 10 µM Y-27632. We used a Cell Lifter (Greiner) to scrape the cells while keeping the cell clumps. One million cells were collected into a 1.5 ml microtube and centrifuged at 200 ×*g* for 3 min at room temperature. We removed the supernatant and tapped the cell pellets. A mixture of 50 µl of GFR Matrigel Phenol Red Free and 50 µl of mTeSR Plus with 10 µM Y-27632 was added and gently mixed with the cells. The cell suspension was aspirated by a 1 ml syringe (TERUMO) with an 18G needle (TERUMO) and stored on ice until transplantation. Mice were anesthetized by 2% isoflurane (VIATRIS). A 1-cm incision was made on the abdominal region, and the epididymal fat pad was carefully pulled out along with the testis. The cell suspension was injected using a syringe and held for 10 sec to avoid backflow. The testis was pushed back to the origin location, and the incision was sutured.

### Statistical analyses

Data are shown as the mean ± standard error of the mean (SEM) in all graphs generated using the Microsoft Excel software program (version 16.66 for Mac; Microsoft). For comparison of two samples, P-values were determined by an unpaired two-tailed Student’s *t*-test using the Microsoft Excel software program (version 16.66 for Mac; Microsoft).

## Acknowledgments

We thank Drs. Ikuo Kawashima (Laboratory of Biomembrane) and Yoichi Tajima (Genome Dynamics Project) for their technical assistance. We also thank all lab members for their helpful discussions.

## Author contributions

I.N. and Y.M. designed the experiments. Y.M. and H.S. generated genome-edited iPSCs and mouse zygotes, respectively. T.O. conducted the co-culture experiments. I.N. and T.O. transplanted iPSCs to mice. T.Tsukimura, T.S., T.Togawa performed the mass spectrometry. I.N. conducted all other experiments. I.N. and Y.M. wrote the manuscript with help from other authors. Y.M. and H.S. supervised the projects.

## Declaration of interests

The authors declare no conflicts of interest in association with the present study.

## Supplemental Information

### Supplemental Experimental Procedures

#### Genome editing in mouse zygotes

The Tokyo Metropolitan Institute of Medical Science Committee on animal research approved the animal experiments of this study (#22-016), and all the experiments were carried out according to the guidelines. The NOD SCID mouse line was purchased from the Charles River Laboratories Japan. For genome editing in mouse zygotes, mixtures of Cas9 protein (New England Biolabs, 30 ng/µl) and sgRNA (5 ng/µl), which was synthesized and purified using the MEGAshortscript T7 Transcription kit and the MEGAclear kit (ThermoFisher) were microinjected into the pronuclei of fertilized eggs prepared by *in vitro* fertilization. The microinjected embryos were cultured in M16 medium supplemented with 50 µM EDTA, and two-cell stage embryos were then transferred into the oviducts of pseudo-pregnant female mice. For genotyping *Gla* of newborn mice, we extracted and purified genomic DNA from pieces of ears. The target site was amplified by PCR using *Gla* Fw2 and *Gla* Rv1 primers (Table S2). The DNA sequences of the PCR products were analyzed by Sanger sequencing using *Gla* Fw1 primer (Table S2). For rapid genotyping of the *Gla*^Δ28^ line that was mainly used in this study, the extracted genomic DNA was amplified by PCR using *Gla* Fw1 and *Gla* Rv3 primers (Table S2) and the products were separated by agarose electrophoresis to determine the genotype by the size of PCR products (249-bp and 221-bp for WT and *Gla*^Δ28^, respectively). The PCR conditions used in this study are summarized in Table S3.

### Preparation of cell lysates and culture supernatants to measure GLA activity

Culture supernatants were collected and centrifuged at 1,180 ×*g* for 3 min. For preparation of cell lysates, cells were detached from culture plates and resuspended in PBS. The cell suspension was centrifuged at 1,180 ×*g* for 3 min to remove the supernatant. The cells were washed with 1 ml PBS and centrifuged again. We added 40 µl of MES pH6.0 with cOmplete Mini to the cells. The cells were homogenized by ultrasonication for 1 sec three times on ice using a Handy Sonic UR21-P (TOMY). The homogenate was centrifuged at 1,180 ×*g* for 5 min at 4°C, and the supernatant was used for the assay.

### Preparation of mouse tissues to measure the GLA activity

Mice were anesthetized by intraperitoneal administration of 100 µl of 2% midazolam (Sandoz), 7.5% of domitor (Zenoaq), and 10% of butorphanol (Meiji Seika Pharma) in PBS per 10 g of mouse body weight. We collected 300 µl of blood from the inferior vena cava and carefully mixed it with 2 µl of heparin sodium (Mochida) and incubated at room temperature for 30 min. The mixture was centrifuged at 800 ×*g* for 5 min at 4°C and the blood plasma was collected. Then, the liver, kidneys, heart, and teratoma were sampled. The organs were washed with PBS and a 200 mg piece of the organ was cut out, and 40 µl of MES pH6.0 (Nacalai Tesque) with cOmplete mini (Sigma) was added. The tissues were homogenized by Ultra Turrax (IKA) and ultrasonic waves generated from a Handy Sonic UR21-P (Tomy). The homogenized samples were centrifuged at 13,000 ×*g* for 5 min at 4 °C and supernatant was collected for the assay.

### Feeder-free culture of human iPSCs

For human iPSCs, this study was approved by the Tokyo Metropolitan Institute of Medical Science Committee on human research (#18-44). The human iPSC line used in this study was generated from a healthy male patient, WTC11 (Kreitzer et al., 2013), using the episomal reprogramming method (Okita et al., 2011). iPSCs were maintained on GFR Matrigel Phenol Red Free (Corning) in mTeSR Plus (Stem Cell Technologies) supplemented with 1% Penicillin-Streptomycin (Pen-Sterp), which was exchanged every other day. When the cell density was low, we added 10 µM Y-27632, a Rho-associated kinase (ROCK) inhibitor, to promote cell survival (Watanabe et al., 2007).

### Differentiation of human iPSCs to fibroblasts and cardiomyocytes

For fibroblast differentiation, we followed a previously reported protocol (Xu et al., 2015). In short, iPSCs were seeded into a Matrigel-coated 12-well plate. When the cells reached 90-100% confluence, the medium was changed to Dulbecco’s modified Eagle’s medium (DMEM) with 10% Fetal Bovine Serum (FBS) (Thermo Fisher) and 1% Pen-Strep (Day 0). The medium was changed every 2 days for 14 days. At Day 15, the cells were detached by 0.25% Trypsin-EDTA and replated in a 6-well plate, and then cultured in DMEM with 10% FBS and 1% Pen-Strep. When the cells reached 100% confluence, they were replated into a new 6-well plate at a ratio of 1:20. After repeating this replating 5 times, the cells were detached and 4×10^4^ cells/well were plated into a 24-well plate using DMEM (high glucose) with 50 µg/ml L-ascorbic acid phosphate Magnesium Salt n-Hydrate (Wako) and 100 ng/ml human recombinant CTGF (Wako). The medium was changed every 3 days for 2 weeks, and the cells differentiated into fibroblasts. For cardiomyocyte differentiation, we followed a previously reported protocol (Lian et al., 2012). In short, 1×10^5^ or 2×10^5^ of iPSCs were plated into a 12-well plate. When the cells reached 80-90% confluence, the medium was changed to RPMI1640 with B27(insulin-) (Thermo Fisher) and 6 or 12 µM CHIR99021 (ChemScene) (Day 0). At Day 1 for 6 µM CHIR99021 culture, or Day 2 for 12 µM CHIR99021 culture, the medium was changed to RPMI1640 with B27(insulin-) and 5 µM IWP-2 (R&D Systems). Two days after the addition of IWP-2, the medium was changed to RPMI1640 with B27(insulin-). Two days later, the medium was then exchanged to RPMI1640 with B27 (Thermo Fisher). The same media was changed every 2 or 3 days to generate beating cardiomyocytes.

Then, we followed a protocol of the metabolic selection of cardiomyocytes using lactate (Tohyama et al., 2013). At Day 15, the cells were detached from the 12-well plate with 0.25% Trypsin-EDTA and resuspended in EB20 medium (Knockout DMEM (Thermo Fisher) containing 10% FBS, 1x GlutaMAX-I (Thermo Fisher), 100 µM MEM NEAA (Thermo Fisher), and 0.1 mM β-mercaptoethanol). The cell suspension was centrifuged at 300 ×*g* for 5 min and supernatant was removed. The cells were resuspended in RPMI1640 with B27 and 10 µM Y-27632, and plated into a Matrigel-coated well of a 6-well plate. At Day 16, the medium was changed to RPMI1640 with B27 without 10 µM Y-27632. At Day 20, the medium was changed to Lactate medium (1M HEPES solution (Gibco) containing 1M lactate (Sigma)). The differentiated cardiomyocytes were selected by changing Lactate medium at Day 22 and 24 too. At Day 26, the medium was changed back to RPMI1640 with B27. The purified cardiomyocytes were used for the assays at Day 28.

## Supplemental Tables

**Table S1.**
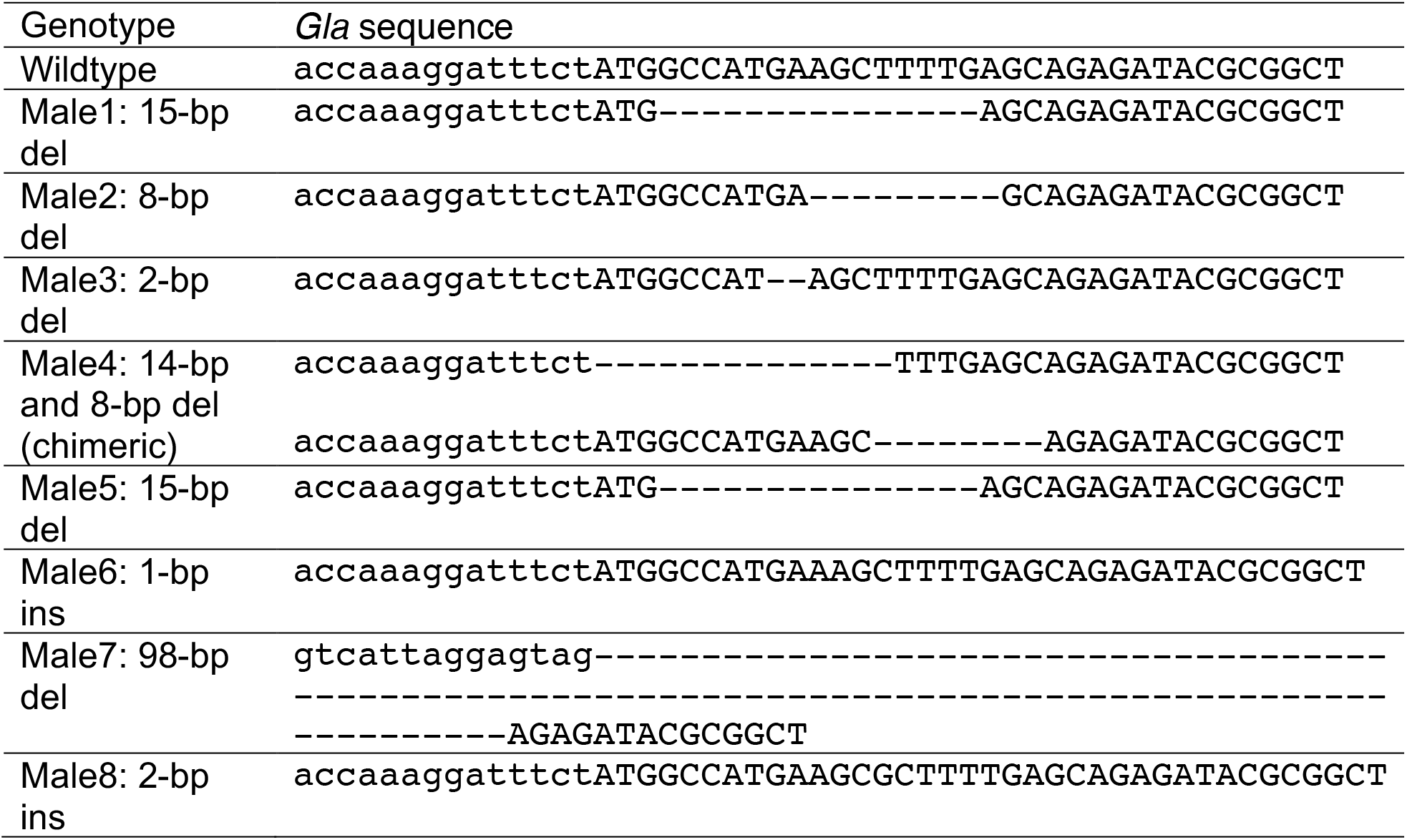
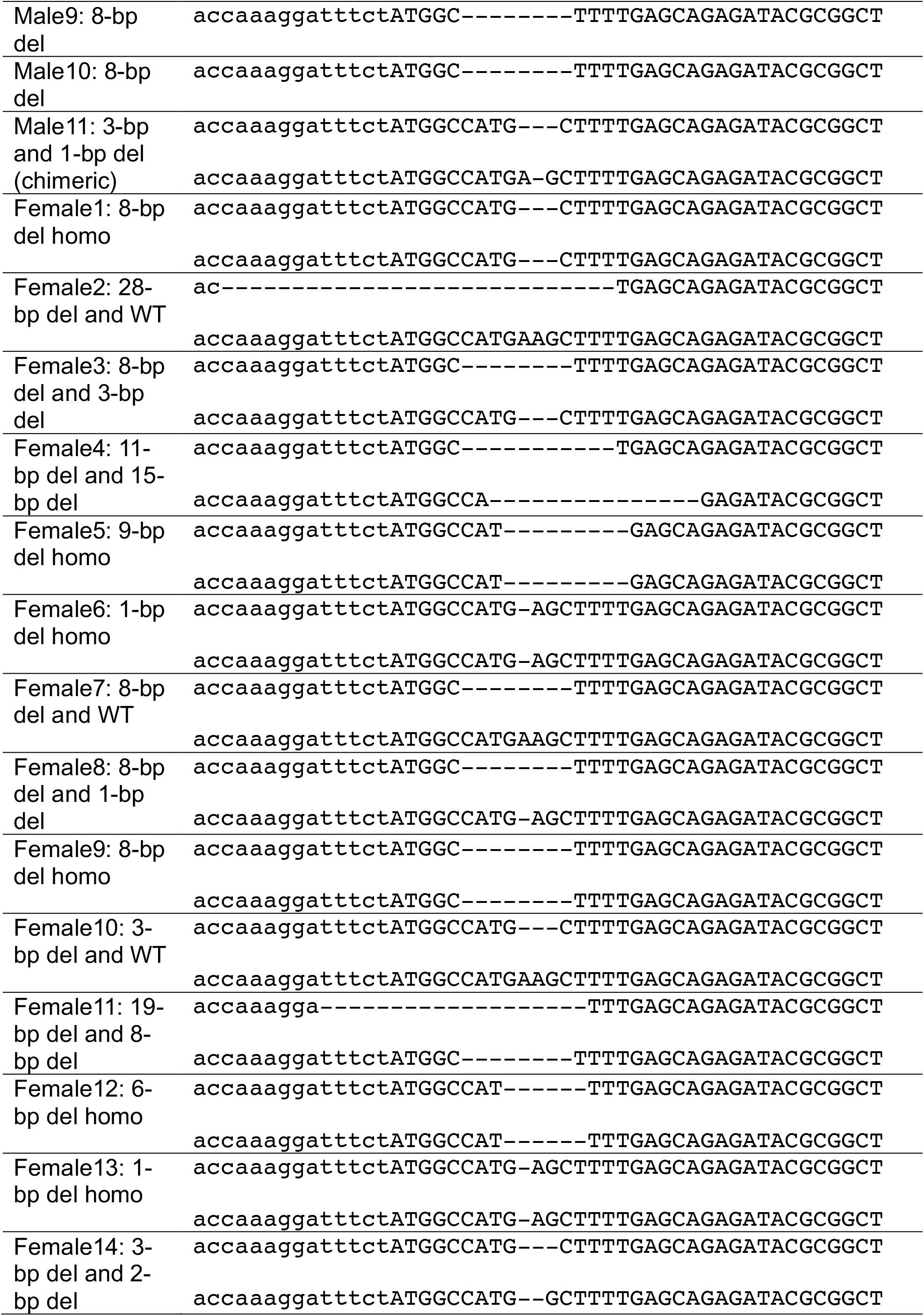
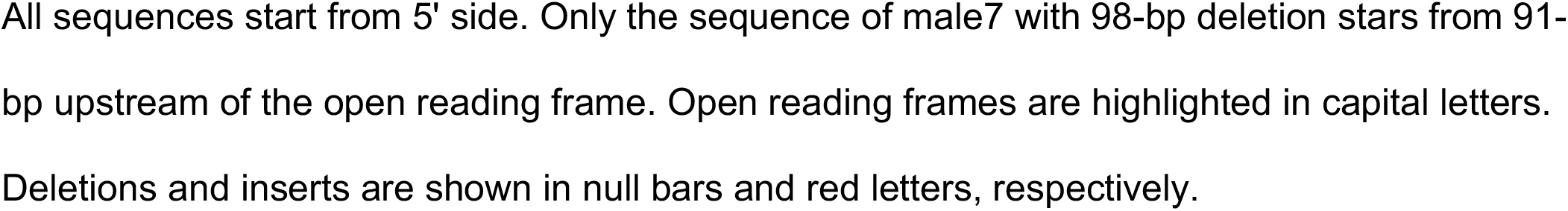
Edited *Gla* alleles of NOD SCID newborn mice. Genotype *Gla* sequence

**Table S2.**
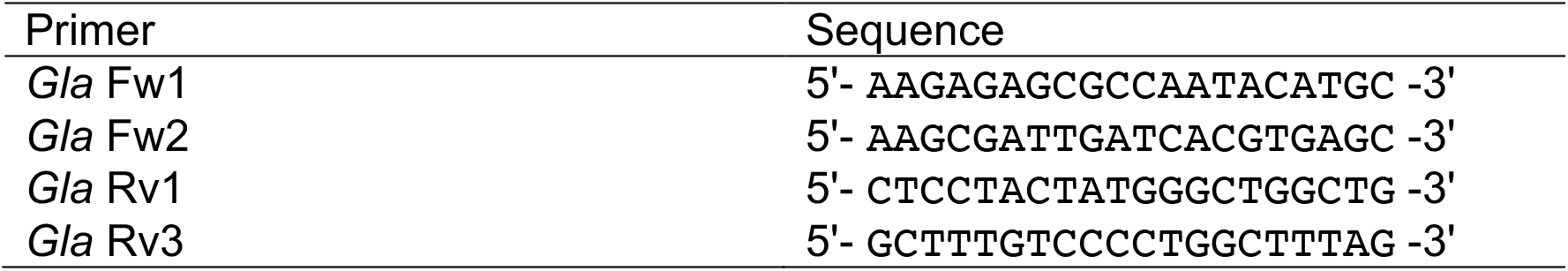
Genotyping primers for mouse *Gla* gene. Primer Sequence

**Table S3.** PCR condotion for mouse *Gla* genotyping.

|  |  |
| --- | --- |
| Step1 | Denaturation : 98°C for 10sec |
| Step2 | Annealing : 52°C for 5sec |
| Step3 | Extension : 68°C for 1sec |
| Step4 | Step1-3 were repeated 34 times |

The PCR conditions for genotyping of mouse *Gla* used in this study is shown. The PCR
reactions took place in a total volume of 25 μl containing 100 ng of purified mouse genomic DNA, 1.5 μl of 10 μM each primer, and 12.5 μl of KOD One PCR Master Mix (TOYOBO). The volume was adjusted to 25 μl with diluted water. The annealing temperature was same for all genotyping primer set used in this study.

## Notes

### Competing Interest Statement

The authors have declared no competing interest.

## References

Benjamin, E.R., Della Valle, M.C., Wu, X., Katz, E., Pruthi, F., Bond, S., Bronfin, B., Williams, H., Yu, J., Bichet, D.G., et al. (2017). The validation of pharmacogenetics for the identification of Fabry patients to be treated with migalastat. Genet. Med. 19, 430–438. https://doi.org/10.1038/gim.2016.122.

Choi, Y.-R., Collins, K.H., Springer, L.E., Pferdehirt, L., Ross, A.K., Wu, C.-L., Moutos, F.T., Harasymowicz, N.S., Brunger, J.M., Pham, C.T.N., et al. (2021). A genome-engineered bioartificial implant for autoregulated anticytokine drug delivery. Sci. Adv. 7, eabj1414. https://doi.org/10.1126/sciadv.abj1414.

Concolino, D., Deodato, F., and Parini, R. (2018). Enzyme replacement therapy: efficacy and limitations. Ital. J. Pediatr. 44, 120. https://doi.org/10.1186/s13052-018-0562-1.

Eng, C.M., Guffon, N., Wilcox, W.R., Germain, D.P., Lee, P., Waldek, S., Caplan, L., Linthorst, G.E., Desnick, R.J., and International Collaborative Fabry Disease Study Group (2001). Safety and efficacy of recombinant human alpha-galactosidase A replacement therapy in Fabry’s disease. N. Engl. J. Med. 345, 9–16. https://doi.org/10.1056/NEJM200107053450102.

Fabry, J. (1898). Ein Beitrag zur Kenntniss der Purpura haemorrhagica nodularis (Purpura papulosa haemorrhagica Hebrae). Arch. f. Dermat. 43, 187–200. https://doi.org/10.1007/BF01986897.

Germain, D.P. (2010). Fabry disease. Orphanet J. Rare Dis. 5, 30. https://doi.org/10.1186/1750-1172-5-30.

Germain, D.P., Hughes, D.A., Nicholls, K., Bichet, D.G., Giugliani, R., Wilcox, W.R., Feliciani, C., Shankar, S.P., Ezgu, F., Amartino, H., et al. (2016). Treatment of Fabry’s Disease with the Pharmacologic Chaperone Migalastat. N. Engl. J. Med. 375, 545–555. https://doi.org/10.1056/NEJMoa1510198.

González, F., Zhu, Z., Shi, Z.-D., Lelli, K., Verma, N., Li, Q.V., and Huangfu, D. (2014). An iCRISPR platform for rapid, multiplexable, and inducible genome editing in human pluripotent stem cells. Cell Stem Cell 15, 215–226. https://doi.org/10.1016/j.stem.2014.05.018.

Hockemeyer, D., Soldner, F., Beard, C., Gao, Q., Mitalipova, M., DeKelver, R.C., Katibah, G.E., Amora, R., Boydston, E.A., Zeitler, B., et al. (2009). Efficient targeting of expressed and silent genes in human ESCs and iPSCs using zinc-finger nucleases. Nat. Biotechnol. 27, 851–857. https://doi.org/10.1038/nbt.1562.

Iriguchi, S., Yasui, Y., Kawai, Y., Arima, S., Kunitomo, M., Sato, T., Ueda, T., Minagawa, A., Mishima, Y., Yanagawa, N., et al. (2021). A clinically applicable and scalable method to regenerate T-cells from iPSCs for off-the-shelf T-cell immunotherapy. Nat. Commun. 12, 430. https://doi.org/10.1038/s41467-020-20658-3.

Jinek, M., Chylinski, K., Fonfara, I., Hauer, M., Doudna, J.A., and Charpentier, E. (2012). A programmable dual-RNA-guided DNA endonuclease in adaptive bacterial immunity. Science 337, 816–821. https://doi.org/10.1126/science.1225829.

Joung, J.K., and Sander, J.D. (2013). TALENs: a widely applicable technology for targeted genome editing. Nat. Rev. Mol. Cell Biol. 14, 49–55. https://doi.org/10.1038/nrm3486.

Kami, D., Yamanami, M., Tsukimura, T., Maeda, H., Togawa, T., Sakuraba, H., and Gojo, S. (2020). Cell Transplantation Combined with Recombinant Collagen Peptides for the Treatment of Fabry Disease. Cell Transplant. 29, 963689720976362. https://doi.org/10.1177/0963689720976362.

Kobayashi, M., Ohashi, T., Kaneshiro, E., Higuchi, T., and Ida, H. (2019). Mutation spectrum of α-Galactosidase gene in Japanese patients with Fabry disease. J. Hum. Genet. 64, 695–699. https://doi.org/10.1038/s10038-019-0599-z.

Mandai, M., Watanabe, A., Kurimoto, Y., Hirami, Y., Morinaga, C., Daimon, T., Fujihara, M., Akimaru, H., Sakai, N., Shibata, Y., et al. (2017). Autologous Induced Stem-Cell-Derived Retinal Cells for Macular Degeneration. N. Engl. J. Med. 376, 1038–1046. https://doi.org/10.1056/NEJMoa1608368.

Mayes, J.S., Scheerer, J.B., Sifers, R.N., and Donaldson, M.L. (1981). Differential assay for lysosomal alpha-galactosidases in human tissues and its application to Fabry’s disease. Clin. Chim. Acta 112, 247–251. https://doi.org/10.1016/0009-8981(81)90384-3.

Medina, D.L., Fraldi, A., Bouche, V., Annunziata, F., Mansueto, G., Spampanato, C., Puri, C., Pignata, A., Martina, J.A., Sardiello, M., et al. (2011). Transcriptional activation of lysosomal exocytosis promotes cellular clearance. Dev. Cell 21, 421–430. https://doi.org/10.1016/j.devcel.2011.07.016.

Medin, J.A., Tudor, M., Simovitch, R., Quirk, J.M., Jacobson, S., Murray, G.J., and Brady, R.O. (1996). Correction in trans for Fabry disease: expression, secretion and uptake of alpha-galactosidase A in patient-derived cells driven by a high-titer recombinant retroviral vector. Proc Natl Acad Sci USA 93, 7917–7922. https://doi.org/10.1073/pnas.93.15.7917.

Miyaoka, Y., Berman, J.R., Cooper, S.B., Mayerl, S.J., Chan, A.H., Zhang, B., Karlin-Neumann, G.A., and Conklin, B.R. (2016). Systematic quantification of HDR and NHEJ reveals effects of locus, nuclease, and cell type on genome-editing. Sci. Rep. 6, 23549. https://doi.org/10.1038/srep23549.

Ohshima, T., Murray, G.J., Swaim, W.D., Longenecker, G., Quirk, J.M., Cardarelli, C.O., Sugimoto, Y., Pastan, I., Gottesman, M.M., Brady, R.O., et al. (1997). alpha-Galactosidase A deficient mice: a model of Fabry disease. Proc Natl Acad Sci USA 94, 2540–2544. https://doi.org/10.1073/pnas.94.6.2540.

Sakuraba, H., Murata-Ohsawa, M., Kawashima, I., Tajima, Y., Kotani, M., Ohshima, T., Chiba, Y., Takashiba, M., Jigami, Y., Fukushige, T., et al. (2006). Comparison of the effects of agalsidase alfa and agalsidase beta on cultured human Fabry fibroblasts and Fabry mice. J. Hum. Genet. 51, 180–188. https://doi.org/10.1007/s10038-005-0342-9.

Schiffmann, R., Kopp, J.B., Austin, H.A., Sabnis, S., Moore, D.F., Weibel, T., Balow, J.E., and Brady, R.O. (2001). Enzyme replacement therapy in Fabry disease: a randomized controlled trial. JAMA 285, 2743–2749. https://doi.org/10.1001/jama.285.21.2743.

Shultz, L.D., Ishikawa, F., and Greiner, D.L. (2007). Humanized mice in translational biomedical research. Nat. Rev. Immunol. 7, 118–130. https://doi.org/10.1038/nri2017.

Sweeley, C.C., and Klionsky, B. (1963). Fabry’s disease: classification as a sphingolipidosis and partial characterization of a novel glycolipid. J. Biol. Chem. 238, 3148–3150. https://doi.org/10.1016/S0021-9258(18)51888-3.

tajima, Y., Kawashima, I., Tsukimura, T., Sugawara, K., Kuroda, M., Suzuki, T., Togawa, T., Chiba, Y., Jigami, Y., Ohno, K., et al. (2009). Use of a modified alpha-N-acetylgalactosaminidase in the development of enzyme replacement therapy for Fabry disease. Am. J. Hum. Genet. 85, 569–580. https://doi.org/10.1016/j.ajhg.2009.09.016.

takahashi, K., Tanabe, K., Ohnuki, M., Narita, M., Ichisaka, T., Tomoda, K., and Yamanaka, S. (2007). Induction of pluripotent stem cells from adult human fibroblasts by defined factors. Cell 131, 861–872. https://doi.org/10.1016/j.cell.2007.11.019.

tsukimura, T., Shiga, T., Saito, K., Ogawa, Y., Sakuraba, H., and Togawa, T. (2021). Does administration of hydroxychloroquine/amiodarone accelerate accumulation of globotriaosylceramide and globotriaosylsphingosine in Fabry mice? Mol. Genet. Metab. Rep. 28, 100773. https://doi.org/10.1016/j.ymgmr.2021.100773.

Urnov, F.D., Rebar, E.J., Holmes, M.C., Zhang, H.S., and Gregory, P.D. (2010). Genome editing with engineered zinc finger nucleases. Nat. Rev. Genet. 11, 636–646. https://doi.org/10.1038/nrg2842.

van Meel, E., Lee, W.-S., Liu, L., Qian, Y., Doray, B., and Kornfeld, S. (2016). Multiple Domains of GlcNAc-1-phosphotransferase Mediate Recognition of Lysosomal Enzymes. J. Biol. Chem. 291, 8295–8307. https://doi.org/10.1074/jbc.M116.714568.

Xu, H., Wang, B., Ono, M., Kagita, A., Fujii, K., Sasakawa, N., Ueda, T., Gee, P., Nishikawa, M., Nomura, M., et al. (2019). Targeted Disruption of HLA Genes via CRISPR-Cas9 Generates iPSCs with Enhanced Immune Compatibility. Cell Stem Cell 24, 566-578.e7. https://doi.org/10.1016/j.stem.2019.02.005.

Yogalingam, G., Bonten, E.J., van de Vlekkert, D., Hu, H., Moshiach, S., Connell, S.A., and d’Azzo, A. (2008). Neuraminidase 1 is a negative regulator of lysosomal exocytosis. Dev. Cell 15, 74–86. https://doi.org/10.1016/j.devcel.2008.05.005.

## Supplemental References

Kreitzer, F.R., Salomonis, N., Sheehan, A., Huang, M., Park, J.S., Spindler, M.J., Lizarraga, P., Weiss, W.A., So, P.-L., and Conklin, B.R. (2013). A robust method to derive functional neural crest cells from human pluripotent stem cells. Am. J. Stem Cells 2, 119–131..

Lian, X., Hsiao, C., Wilson, G., Zhu, K., Hazeltine, L.B., Azarin, S.M., Raval, K.K., Zhang, J., Kamp, T.J., and Palecek, S.P. (2012). Robust cardiomyocyte differentiation from human pluripotent stem cells via temporal modulation of canonical Wnt signaling. Proc Natl Acad Sci USA 109, E1848–57. https://doi.org/10.1073/pnas.1200250109.

Okita, K., Matsumura, Y., Sato, Y., Okada, A., Morizane, A., Okamoto, S., Hong, H., Nakagawa, M., Tanabe, K., Tezuka, K., et al. (2011). A more efficient method to generate integration-free human iPS cells. Nat. Methods 8, 409–412. https://doi.org/10.1038/nmeth.1591.

tohyama, S., Hattori, F., Sano, M., Hishiki, T., Nagahata, Y., Matsuura, T., Hashimoto, H., Suzuki, T., Yamashita, H., Satoh, Y., et al. (2013). Distinct metabolic flow enables large-scale purification of mouse and human pluripotent stem cell-derived cardiomyocytes. Cell Stem Cell 12, 127–137. https://doi.org/10.1016/j.stem.2012.09.013.

Watanabe, K., Ueno, M., Kamiya, D., Nishiyama, A., Matsumura, M., Wataya, T., Takahashi, J.B., Nishikawa, S., Nishikawa, S., Muguruma, K., et al. (2007). A ROCK inhibitor permits survival of dissociated human embryonic stem cells. Nat. Biotechnol. 25, 681–686. https://doi.org/10.1038/nbt1310.

Xu, R., Taskin, M.B., Rubert, M., Seliktar, D., Besenbacher, F., and Chen, M. (2015). hiPS-MSCs differentiation towards fibroblasts on a 3D ECM mimicking scaffold. Sci. Rep. 5, 8480. https://doi.org/10.1038/srep08480.

